# Targeting extracellular vesicle delivery to the lungs by microgel encapsulation

**DOI:** 10.1101/2022.09.09.507125

**Authors:** Nicholas D. Cober, Katelynn Rowe, Yupu Deng, Ainara Benavente-Babace, David W. Courtman, Michel Godin, Duncan J. Stewart

**Author notes:** **Corresponding author**: Duncan J Stewart., Executive Vice-President Research, The Ottawa Hospital; CEO & Scientific Director, Ottawa Hospital Research Institute; The Evelyne & Rowell Laishley Chair and Professor, Department of Medicine, University of Ottawa, 501 Smyth Road, Box/C.P. 511, Ottawa, ON, K1H 8L6.

## Abstract

Extracellular vesicles (EVs) secreted by stem and progenitor cells have significant potential as cell-free ‘cellular’ therapeutics. Yet, small EVs (<200 nm) are rapidly cleared after systemic administration, mainly by the liver, presenting challenges targeting EVs to a specific organ or tissue. Microencapsulation using natural nano-porous hydrogels (microgels) has been shown to enhance engraftment and increase the survival of transplanted cells. We sought to encapsulate EVs within microgels to target their delivery to the lung by virtue of their size-based retention within the pulmonary microcirculation. Mesenchymal stromal cell (MSC) derived EVs were labelled with the lipophilic dye (DiR) and encapsulated within agarose-gelatin microgels. Endothelial cells and bone marrow derived macrophages were able to take up EVs encapsulated in microgels *in vitro*, but less efficiently than the uptake of free EVs. Following intrajugular administration, microgel encapsulated EVs were selectively retained within the lungs for 72 hours, while free EVs were rapidly cleared by the liver. Furthermore, microgel loaded EVs demonstrated greater uptake by lung cells, in particular CD45+ immune cells, as assessed by flow cytometry compared to free EVs. Microencapsulation of EVs may be a novel tool for enhancing targeted delivery of EVs for future therapeutic applications.

## Introduction

There is increasing evidence that extracellular vesicles (EVs) play an essential role in cell-to-cell communication.^1^ EVs are nano-sized membrane bound particles loaded with proteins, mRNAs, and miRNAs from their host cell which provide important signaling cues to their recipient cell.^1^ EVs can be categorized based on their size; small EVs are <150nm in size likely contain the exosome fraction of endosomal origin, whereas EVs between 150-1000nm contain primarily microvesicles.^2^ Therefore, there is considerable interest in developing stem cell derived EVs, particularly small EVs, as a cell-free ‘cellular’ therapeutic product.^3^ Mesenchymal stromal cells (MSCs) represent a multi-faceted therapeutic cell with important angiogenic and immunomodulatory potential and have been reported to have therapeutic effects in a number of models of cardiovascular disease.^4^ The actions of MSCs are thought to be mediated largely by paracrine mechanisms, in particular related to the release of therapeutic EVs.^3^ In fact, MSC-EVs have been studied as a therapy for a variety of cardiovascular diseases including pulmonary arterial hypertension (PAH)^5–7^, acute myocardial infarction^7,8^, acute lung injury^9^, and bronchopulmonary dysplasia^10^.

EVs have a number of potential advantages over intact cells. EVs have the potential to mirror the benefits of cell-based therapies without some of the risks associated with delivery of intact cells, including tumorigenicity, immunoreactivity, product shelf life, and viability.^11^ As EVs are cell by-products they are unable to replicate and are therefore not tumorigenic. Furthermore, the transplantation of allogenic cells presents a risk of immunoreactivity from foreign antigens, whereas EVs derived from MSCs generally lack components of the major histocompatibility complex.^12^ Finally, unlike intact cells where efficacy is dependent on cell viability that can be lost during storage,^13^ EVs possess good long term stability at −80°C^14^ and do not have post thaw viability requirements that may hinder their use.

However, unlike intact cells which can be used to target the lung through size-based filtration in the distal lung arteriolar bed, small EVs, in the nanometer range, would not be expected to lodge in the precapillary arteriolar bed and are rapidly cleared from the circulation by the liver and spleen^15,16^, limiting their therapeutic potential. Indeed, Gupta et al. have shown in a pharmacokinetic study that EVs have a very short half-life *in vivo* and are cleared from the blood within minutes of delivery.^17^ Furthermore biodistribution may be impacted by cell origin; while EVs from many cell types largely accumulate in the liver and spleen,^15^ EVs derived from breast cancer cells may preferentially target the lung,^18^ as well as by modifications, such as glycolsylation,^19^ or genetic modification.^20^

Biomaterials are commonly used to create cell compatible microenvironments, promoting cell viability and facilitating transplantation,^21^ and can be engineered to target distribution to a particular organ. Microencapsulation can enhance local organ retention of intact cells^22^ and has been shown to promote therapeutic outcomes in experimental models of acute myocardial infarction.^23^ More recently, bulk EV encapsulation has been used to create large regenerative tissue patches for cardiac repair.^24^ We hypothesized that microencapsulation of EVs in nano-porous hydrogels (microgels), in a manner suitable for intravascular delivery, would promote local EV retention within the lung, facilitating increased cellular uptake. We now report markedly enhanced retention of EV-loaded microgels within the lungs as compared to free EVs, with enhanced local cellular uptake, supporting the use of microencapsulation as a novel strategy to efficiently target EVs to the lung.

## Methods

### Mesenchymal stromal cell culture and extracellular vesicle isolation

Sprague Dawley rat bone marrow mesenchymal stromal cells (MSCs) were isolated from rat bone marrow as previously described^25^ or acquired from Cyagen (Cyagen Bioscience, Santa Clara, CA, USA). Isolated rat bone marrow MSCs were cultured in alpha MEM (ThermoFisher Scientific, Burlington ON, Canada) with 10% fetal bovine serum (ThermoFisher Scientific, Burlington ON, Canada) and 1% pencilin/streptomycin (Life Technologies, Carlsbad, CA, USA), Cyagen sourced MSCs were cultured in StemXVivo mesenchymal stem cell expansion media (R&D systems, Toronto, ON, Canada) inside tissue culture incubators at 37°C with 5% CO_2_, passaging as necessary.

### Extracellular vesicle isolation and labelling

MSCs between passage 3 and 5 were used for EV isolations. For EV isolation, MSCs were cultured to 80-90% confluency, washed 2x with PBS and changed to serum free alpha-MEM (Invitrogen, Waltham, MA, USA) with 1% pencilin-streptomycin. MSC-conditioned media was collected after purified using sequential ultracentrifugation and tangential flow filtration (TFF) (KrosFlow KR2i, Repligen, Waltham, MA, USA). In brief, conditioned media (CM) was spun at 2500g for 10min at 4C, the supernatant was then spun at 20,000g for 20min at 4C. The resulting conditioned media was processed by TFF to remove small protein and particles with a 500,000 MWCO MidGee hoop ultrafiltration cartridge (GE Lifesciences, Vancouver, BC, Canada).

Diafiltration was performed with sterile filtered PBS and concentrated sample was collected. To further purify small EVs, conditioned media was spun at 100,000g for 30min at 4°C and pellets were resuspended in sterile PBS. For DiR (ThermoFisher Scientific, Burlington, ON, Canada) and DiD (ThermoFisher Scientific, Burlington, ON, Canada) labelling, concentrated EVs from the TFF were stained with DiR or DiD at 0.5 ug/ml for 15min than spun at 100,000g for 30min at 4°C to wash unbound dye and pellets were resuspended in PBS. For pkh26 (Sigma Aldrich, Oakville, ON, Canada) labelling, concentrated EV fraction following TFF was spun at 100,000g for 30min at 4°C, and pellets were resuspended in dilutant C. pkh26 (0.002mM; Sigma Aldrich, Oakville, ON, Canada) was added for 15 min following which sample was spun at 100,000g for 30min at 4°C to concentrate and wash the EV fraction. Labelled EVs were resuspended in PBS, aliquoted and stored at −80°C until use.

### Extracellular vesicle characterization

MSC derived small EVs were characterized for protein concentration using the bicinchoninic acid (BCA) assay (Sigma Aldrich, Oakville, ON, Canada). Nanoparticle tracking analysis was performed using ZetaView (Particle Metrix, Mebane, NC, USA) to determine particle number and size distribution. Western blot was performed to confirm presence of canonical EV protein markers CD63, CD9, TSG101, or negative selection marker GM-130. Samples were lysed in 1x RIPA buffer (Millipore, CA, USA) + Halt protease inhibitors (ThermoFisher Scientific, Burlington, ON, Canada) with sonication and vortexing for protein quantification, as above. Loading buffer was added and samples were heated at 70°C for 10min. Samples (10-15ug protein/lane) were run by electrophoresis on pre-cast SDS-Page Stain Free Gels (Bio-Rad, Hercules, CA, USA). Separated proteins were transferred to low fluorescence PVDF membranes using the TransBlot Turbo (Bio-Rad, Hercules, CA, USA), and blocked in TBS-T (TBS with 0.1% Tween 20) with 5 % non-fat dry milk for 1h at room temperature. Membranes were washed 5x times with TBS-T and stained overnight at 4°C with gentle agitation with primary antibodies: mouse anti-rat CD63 (1:200 dilution, BD Biosciences, Mississauga, ON, Canada), rabbit anti-rat CD9 (1:500 dilution, ThermoFisher Scientific, Burlington, ON, Canada), rabbit anti-rat TSG101 (1:500 dilution, ThermoFisher Scientific, Burlington, ON, Canada), or rabbit anti-mouse GM-130 (1:500 dilution, ThermoFisher Scientific, Burlington, ON, Canada). After washing membranes 5x in TBS-T membranes were incubated with appropriate horseradish peroxidase (HRP)-conjugated secondary antibodies (1:5000, Jackson ImmunoResearch West Grove, PA, USA) for 1h at room temperature. Clarity Western ECL (Bio-Rad, Hercules, CA, USA) was used to detect HRP signal imaging with ChemiDoc MP imaging system (Bio-Rad Hercules, CA, USA). Images were analyzed with Image Lab v6 (Bio-Rad, Hercules, CA, USA). Data shown in **Supplemental Figure 1**.

### HUVEC uptake experiments

Human umbilical vein endothelial cells (HUVEC) were cultured in endothelial cell growth media with 5% FBS, 1% pen/strep and supplements (ScienCell, Carlsbad, CA, USA), in a tissue culture incubator at 37°C with 5% CO_2_, passaging as needed. HUVEC between passages 4 and 6 were used for experiments. HUVEC were seeded at 100,000 cells/ well in a 6 well plate (Corning, Corning, NY, USA), after 24h DiR or DiD labelled free EVs were added at 1ug or 3ug of EV protein. After 24h of uptake, cells were washed with PBS, lifted with trypLE (Life Technologies, Carlsbad, CA, USA), and resuspended in **PEB**: **P**BS + 2mM **E**DTA (ThermoFisher Scientific, Burlington, ON, Canada) + 0.5% **b**ovine serum albumin (BSA, Wisent Bioproducts, Saint-Jean-Baptiste, QC, Canada). Cells were counted, washed, and resuspended in staining buffer containing: 1:100 dilution V450 mouse anti-human CD31 (BD Biosciences, Mississauga, ON, Canada), 1:10 dilution PE mouse anti-human CD34 (BD Biosciences, Mississauga, ON, Canada), for 30min at 4°C in the dark. Appropriate unstained, fluorescent minus one (FMOs), and compensation controls were prepared. Following staining cells were washed with PEB and loaded into 5ml flow tube (Corning, Corning, NY, USA) for flow analysis on the Attune NXT (ThermoFisher Scientific, Burlington, ON, Canada), or the AMNIS ImageStream X (Luminex, Austin, TX, USA) for intracellular visualization. Subsequent analysis was performed in FlowJo version 10 (FlowJo LLC, Ashland, OR, USA).

### Bone marrow derived macrophage isolation, characterization, and uptake

Rat bone marrow derived macrophages were derived, as adapted from Souza-Moreira et al.^26^. Rat bone marrow was isolated, mononuclear cells were counted, and frozen in 90% heat inactivated FBS (ThermoFisher Scientific, Burlington, ON, Canada) + 10% DMSO (Sigma Aldrich, Oakville, ON, Canada). Rat bone marrow mononuclear cells were cultured at 4-6 million in a 10cm dish (Falcon) using differentiation media containing: DMEM (ThermoFisher Scientific, Burlington, ON, Canada), 20% HI FBS, 1% Pen/strep, 1% Glutamax (ThermoFisher Scientific, Burlington, ON, Canada), and 20 ng/ml macrophage colony stimulating factor (MCSF, Sigma Aldrich, Oakville, ON, Canada) in a tissue culture incubator at 37°C, 5% CO_2_. Additional media was added after 3-4 days. After 6-7 days macrophages were lifted using cold PBS and scraping than counted. Bone marrow derived macrophages were plated at 1 million per well, in a 6 well plate in DMEM + 10% HI FBS, 1%Pen-strep, 1%glutamax with or without LPS (50 ng/ml, Sigma Aldrich, Oakville, ON, Canada) + interferon gamma (IFNγ, 5 ng/ml PeproTech Inc., Cranberry, NJ, USA) or interleukin 4 (IL4, 10 ng/ml, PeproTech Inc., Cranberry, NJ, USA) stimulation. Macrophages were characterized for presence and absence of canonical markers using flow cytometry PE-Cy5 mouse anti-rat CD45 (BD Biosciences, Mississauga, ON, Canada), BV786 mouse anti-rat CD11b/c (BD Biosciences, Mississauga, ON, Canada), and PE mouse anti-rat CD34 (Novus Biologicals, Littleton, CO, USA). (**Supplemental Figure 2**). For uptake experiments macrophages were cultured, lifted, and plated for 24h of stimulation with or without LPS+IFNg. Subsequently, macrophages were washed with PBS, and fresh media was added with or without IL4, and DiR labelled EVs at 1ug of EV protein. After 24h macrophages were lifted, washed, counted, and stained. Staining was performed with 1:200 dilution eFluor 450 mouse anti-rat CD45 (eBioscience, San Diego, CA, USA), and FITC mouse anti-rat CD86 (eBioscience, San Diego, CA, USA), for 30min at 4°C in the dark. Following which cells were washed and analyzed by flow cytometry as previously described.

### Microfluidic EV encapsulation

Extracellular vesicles were encapsulated using a novel microfluidic device designed for cell and small particle encapsulation, in a similar manner as described for cell encapsulation^27,28^. In brief, a known quantity of EV protein was mixed with 1% ultra-low gelling temperature agarose (Sigma Aldrich, Oakville, ON, Canada), 1% gelatin (Sigma Aldrich, Oakville, ON, Canada) and kept at 37°C during encapsulation. Following microfluidic encapsulation, microgels were kept at 4°C to ensure complete gelation, and 3x PBS washes with 1000g for 5min centrifugation spins to remove the residual oil. Microgels were counted using the hemocytometer and used for experiments based on initial EV protein incorporated into total batch of microgels. Microgels were stored at 4°C for up to 24h prior to *in vivo* injection. For fluorescent visualization gelatin – Oregon green 488 (ThermoFisher Scientific, Burlington, ON, Canada) was added to capsules at a diluted concentration (0.5%). Fluorescently labeled EV loaded and empty microgels were visualized with both a Zeiss M2 imager epi-fluorescent and a Zeiss LSM900 confocal microscope.

### MCT Model of PAH and biodistribution

All animal experiments were approved by the University of Ottawa Animal Care Committee. As previously described,^29^ male Sprague Dawley (SD) rats (200-250g) were injected with monocrotaline (MCT) (60 mg/kg, Sigma Aldrich, Oakville, ON, Canada) by intra peritoneal (i.p.) injection seven days before EV injections. For EV injections, animals were anaesthetized by isoflurane inhalation for jugular vein cut down and cannulation. After wound closure, topical bupivacaine was applied, and administered twice daily for one day following surgery. Buprenorphine (s.c. 0.03 mg/kg, Ceva Santé Animale, Libourne, France) was also administered 1 hour prior to surgery for pain management. Treatments were delivered by intra jugular vein (i.j.) injection using PBS (ThermoFisher Scientific, Burlington, ON, Canada) vehicle. For biodistribution experiments 20ug of DiR labeled free or encapsulated EVs were tracked at 4h, 24h, and 72h post injection by ex vivo imaging with the IVIS spectrum (Perkin Elmer, Waltham, MA, USA). Animals were euthanized, organs (lungs, liver, spleen, kidney, and heart) were isolated and fluorescent images were acquired. Region of interest selection for average radiant efficiency were selected using Living Image Software v 3.2 (Perkin Elmer, Waltham, MA, USA) intensity and fluorescent background was subtracted using vehicle controls.

### Lung digestion and flow analysis

As described, EV therapies were injected 7 days following MCT injection. Lung digestion was performed as previously described^30,31^. 24h post therapy injection animals were euthanized, and lungs were flushed with PBS perfusion through the right heart. Lungs were extracted, manually diced and placed in OCTO-Macs dissociation tubes (Miltenyi Biotec, Bergisch Gladbach, Germany) with digestion buffer [collagenase tvpe 1 (Worthington Biochem., Lakewood, NJ, USA neutral protease (Worthington Biochem., Lakewood, NJ, USA), DNAase I (Sigma Aldrich, Oakville, ON, Canada) in Hank’s Balanced Salt Solution (HBSS, ThermoFisher Scientific, Burlington, ON, Canada)]. Dissociation was performed using OCTOmacs dissociator (Miltenyi Biotec, Bergisch Gladbach, Germany). Following which, digested lungs were filtered through 70um filters (Corning, Corning, NY, USA) and washed with PEB. Red blood cell lysis was performed with 1x RBC lysis buffer (eBioscience, San Diego, CA, USA) for 3min at room temperature, followed by washing with PEB. Cells were counted in PEB and ~1 million cells were taken for staining and appropriate controls. Cells were stained with first Live/Dead Fixable yellow (ThermoFisher Scientific, Burlington, ON, Canada), and then 1:200 dilution of Alexa Fluor 488 mouse anti-rat CD45 (BioLegend, San Diego, CA, USA), PE mouse anti-rat CD31 (BD biosciences, Mississauga, ON, Canada) for 30 min at 4°C in the dark. Cells were washed with PEB and analyzed by flow cytometry on the Attune Nxt (ThermoFisher Scientific, Burlington, ON, Canada), with subsequent analysis performed in FlowJo v10 (FlowJo LLC, Ashland, OR, USA). Gating strategy shown in **Supplemental Figure 3**.

### Statistical Analysis

All data are presented as means ± SEM. Differences between groups were analyzed by one-way with Tukey’s post-hoc test for multiple comparisons or two-way analysis of variance (ANOVA) with Sidak’s post-hoc test for multiple comparisons. An adjusted p-value of p<0.05 was considered significant. All statistical analysis was performed with Graph Pad Prism 8.0 (Graph Pad, San Diego, CA, USA).

## Results

### Differential uptake of extracellular vesicles by cell type

Small EVs are known to play a role in cellular communication in part mediated by uptake by other cells. Small MSC-derived EVs labelled with the near infrared lipophilic dye DiR were incubated with HUVEC or bone marrow-derived macrophages *in vitro*. Less than a third of HUVECs (29 ± 2%) were observed to take up DiR labelled EVs over 24h at 1ug of protein (**Figure 1A-C**), although this increased with higher protein concentration (62 ± 3% at 3ug). Visualization with the AMNIS image stream (**Figure 1B**) and subsequently confocal imaging (**Supplemental Figure 4**) demonstrated the labelled EVs were internalized, not only bound to the cell surface. Compared to HUVECs, a greater proportion of untreated macrophages (MØ) were readily able to take up EVs (75 ± 4% at 1ug) (**Figure 1D-E**) which had no discernable effect on MØ morphology (**Supplemental Figure 5**). Pretreatment with IL4 to induce a M2 alternative activation state, but not LPS which promotes an M1 phenotype, significantly reduced the uptake of MSC EVs (p=0.03) (**Figure 1D-E**).

**Figure 1:**
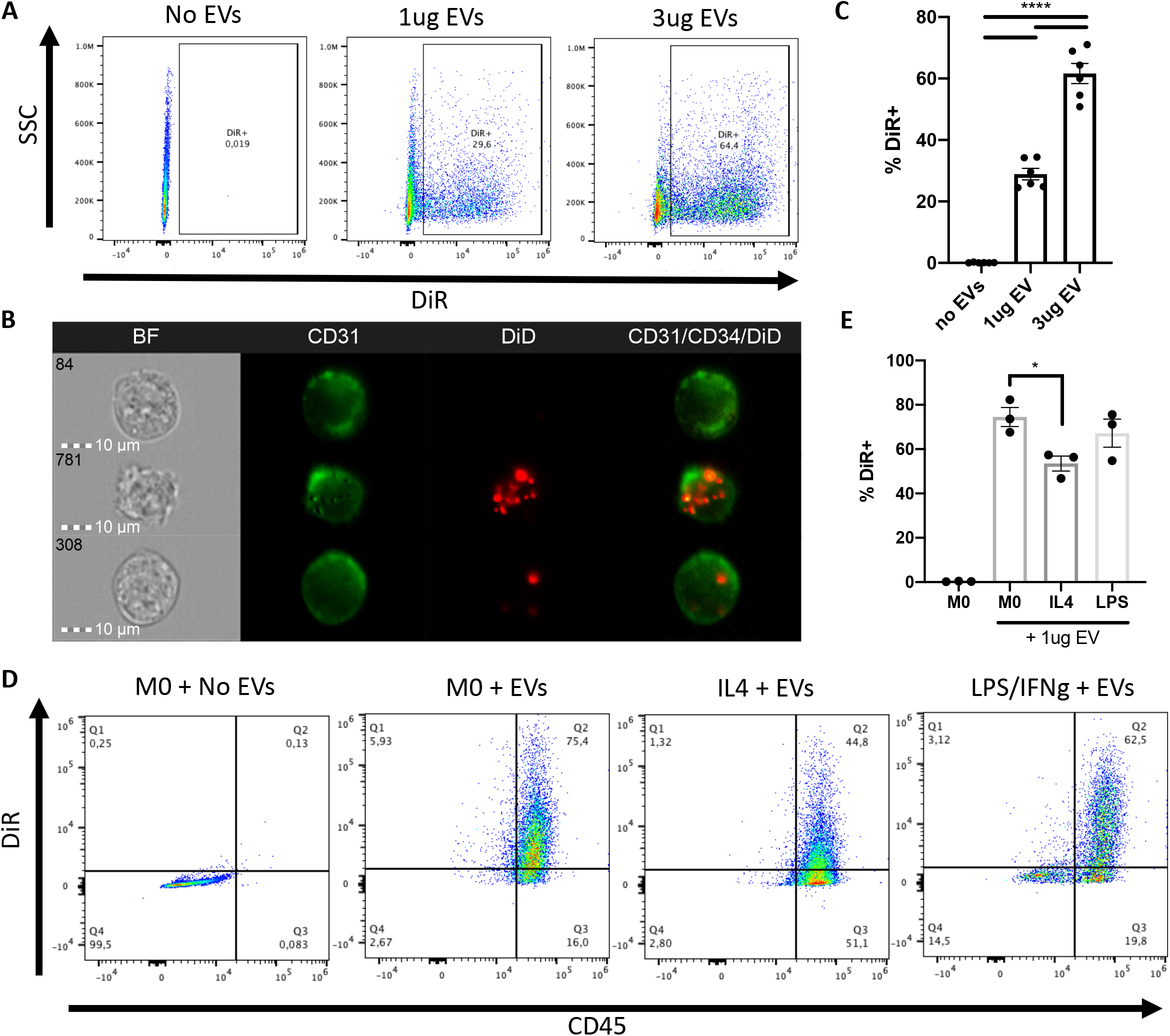
Endothelial cells and macrophages have differential MSC-EV uptake. HUVEC and macrophages were exposed to DiR labelled MSC-EVs *in vitro*. EV uptake by HUVEC was characterized after 24h by (A) flow cytometry and (B) AMNIS analysis revealing the internalization of EVs by a limited population of HUVEC, as quantified (C). Unstimulated macrophages (MØ), IL4 stimulated and LPS/IFNg stimulated macrophages readily took up macrophages as shown in representative flow plots (D) and quantified (E). Data represents mean ± SEM.

### Encapsulation of EVs in microgels

Nanoporous hydrogels have been previously used to encapsulate single cells forming a “microgel” niche to protect cells for *in vivo* delivery^23^. A microfluidic approach^27,28^ was employed to encapsulate EVs labelled with pkh26 in 1% agarose-1% gelatin producing microgels of uniform size (47 ± 0.4 μm) (**Figure 2A-C**). Labelled EVs were detected within microgels evenly dispersed throughout the spherical microgel as observed using confocal imaging z-stacks (**Figure 2D**). Furthermore, *in vitro* encapsulated EVs were taken up by both endothelial cells (9.7 ± 1%) and MØ (64 ± 4%) over 24 hours (**Figure 2E-F**), and again, uptake of EVs was greater for MØ. Integrated fluorescence density per cell was calculated as a measure of relative EV uptake. Interestingly, there was no difference in the proportion of MØ taking up free versus encapsulated EVs (p=0.27), however individual MØ took up significantly more free EVs than encapsulated EVs evidenced by higher fluorescent intensity (p=0.003). This demonstrates that microencapsulation delays the uptake of EVs thereby potentially providing a system for controlled delivery of EVs to a target tissue. Interestingly, while the proportion of HUVECs taking up encapsulated EVs was lower than for free EVs, the amount of uptake per cell was if anything higher (**Figure 2F**), again illustrating important differences in the uptake of EVs between cell types.

**Figure 2:**
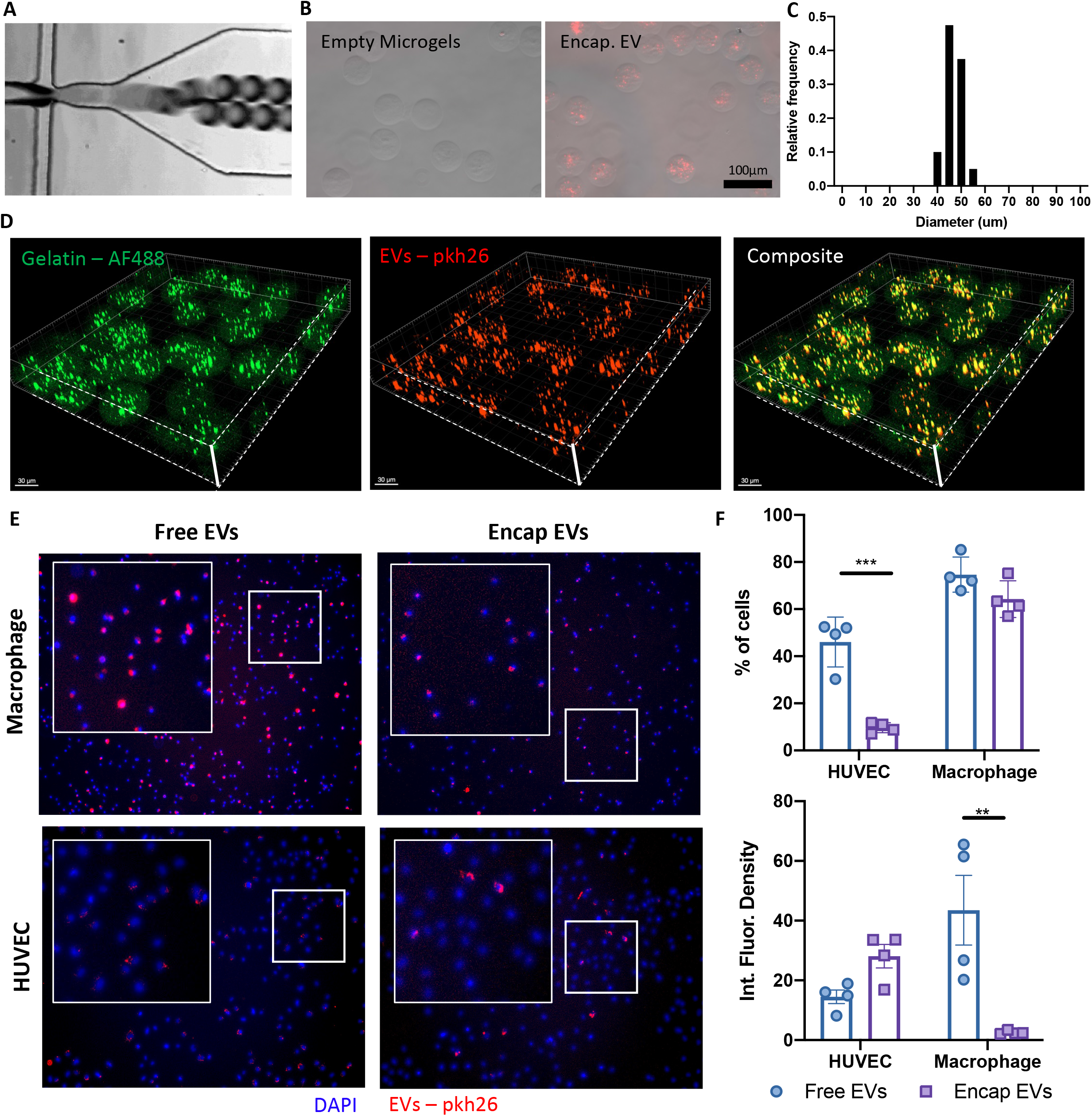
Microencapsulation of MSC-EVs within nanoporous hydrogels. (A) Microfluidic encapsulation was performed with 1%agarose-1%gelatin hydrogels using an oil immersion process to form a homogenous population of EV-loaded microgels. Pkh26 labelled encapsulated EVs were visualized by fluorescent microscopy (B) and microgel diameter quantified (C). Confocal z-stacks demonstrate the loading of EVs throughout the hydrogels (D). Uptake of encapsulated EVs was compared to free EVs after 24h by immunofluorescent imaging (E) and quantification (F) demonstrated the reduced rate of EV uptake compared to free EVs. Data represents mean ± SEM.

### Microencapsulated EVs enhance local lung specific delivery

To evaluate the impact of encapsulation on EV retention, biodistribution experiments were performed using DiR labeled EVs which has a near-infrared fluorescent signal (ex. 750nm; em. 780nm) where tissues have improved optical transparency for *in vivo* imaging and limited auto-fluorescent. Free or encapsulated EVs and vehicle controls were injected through the jugular vein of SD rats 7 days after administration of MCT (**Figure 3A**). At 24h there was a clear difference in biodistribution between free EVs which were predominately taken up by the liver with lung representing only 7.5 ± 0.4% of the measured fluorescent signal, whereas EV-loaded microgels were efficiently retained within the lung (**Figure 3B-C**) representing 81 ± 12 % of the signal (p=0.0001). In contrast, EV-loaded microgels show minimal accumulation within the liver at 24h (10 ± 3%) compared to 74 ± 2% for free EVs (p=0.0001). Accumulation of free and encapsulated EVs was not statistically significant in any other organ examined, including spleen, kidney, and heart (**Supplemental Figure 6**). Surprisingly, the fluorescent signal measured in the lungs of encapsulated EVs actually increased overtime (**Figure 3D-E**). It is possible that at early time points the low fluorescent signal is an artefact caused by quenching of the fluorescent signal of EVs tightly packed inside microgels^32^ and that the quenching is reduced as EVs are progressively released over time and become distributed over a greater area. An alternative explanation would be that EVs are taken up over time from the circulation; however, this is unlikely since the microgels are too large to pass through the lung and are efficiently retained in the lung distal arteriolar bed on first passage. Furthermore, empty 1% agarose-1% gelatin microgels are largely cleared over the first 72h (**Supplemental Figure 7**). However, the lipophilic dye will persist in the membrane of cells after uptake of EVs and this likely accounts for the relatively stable signal intensity from 4 to 72h after delivery of free EVs in the liver, spleen, kidney, and heart (**Supplemental Figure 6**). These findings demonstrate that microencapsulation of small EVs can lead to significant improvement in local lung delivery compared to systemic injection of free EVs.

**Figure 3:**
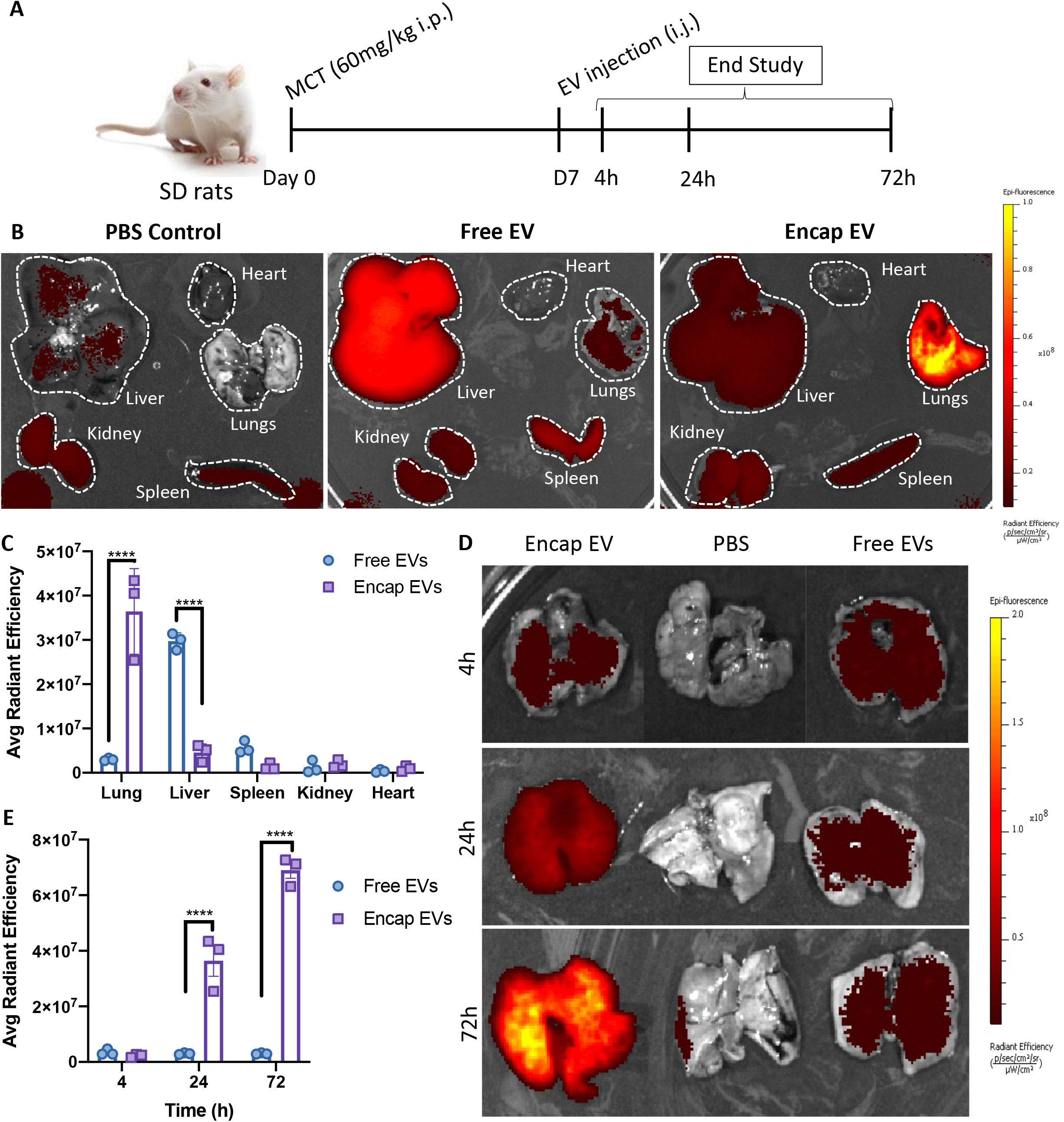
EV loaded microgels were retained within the lungs. (A) Free and encapsulated DiR labelled EVs were administered 7 days after monocrotaline (MCT) which induces a model of pulmonary arterial hypertension. (B) Representative images from IVIS analysis demonstrate the level of background fluorescence observed at 24h in PBS controls compared to free and encapsulated EVs. (C) Quantification of average radiant efficiency at 24h from the lungs, liver, spleen, kidney, and heart demonstrate the significant increase in lung specific retention by encapsulated EVs while free EVs were rapidly cleared to the liver. (D) Representative lung samples from 4h, 24h and 72h time points demonstrate the low level of free EV retention within the lungs, which is further quantified (E). Data represents mean ± SEM.

### Encapsulated EVs were taken up by lung resident immune cells *in vivo*

While the distribution kinetics of free EVs have been studied following systemic administration, little is known about the cellular uptake of these membrane-bound particles in the target organs. We performed lung digestion combined with flow cytometric analysis following the *in vivo* administration of DiR labelled encapsulated or free EVs to identify which cells were interacting with the administered EVs in the lung. The digestion and isolation procedure was biased towards immune cells, with a large proportion of cells being CD45+ across all treatment groups (**Figure 4A-B**). Intravenous injection of microgels, with or without, EVs resulted in an increase in CD45+ cell recruitment to the lungs compared to healthy controls (**Figure 4B**), consistent with a proinflammatory effect of agarose-based hydrogel. Histological data corroborated this finding demonstrating cellular influx surrounding microgels (**Supplemental Figure 7**). No statistical difference was observed between number of endothelial cells (CD45-CD31+), or other cells (CD45-CD31-) (e.g. fibroblasts, epithelial) in control lungs and those from animals receiving microgels (**Figure 4B**). Interestingly, at 24h, only animals treated with EV-loaded microgels showed statistically significant accumulation of DiR+ EVs within both the CD45+CD31- and CD45+CD31lo immune cell populations compared to all other treatment groups (**Figure 4C-D**). Despite the relative low yield of endothelial cells (CD45-CD31+) from the digestion, EV uptake was seen only in animals receiving EV-loaded microgels. No differences were observed in the CD45-CD31-cells uptake of DiR+ EVs between all the treatment groups (**Supplemental Figure 8**). Therefore, these findings demonstrate that EVs are predominately taken up by immune cells *in vivo*, and encapsulation enhances EV uptake.

**Figure 4:**
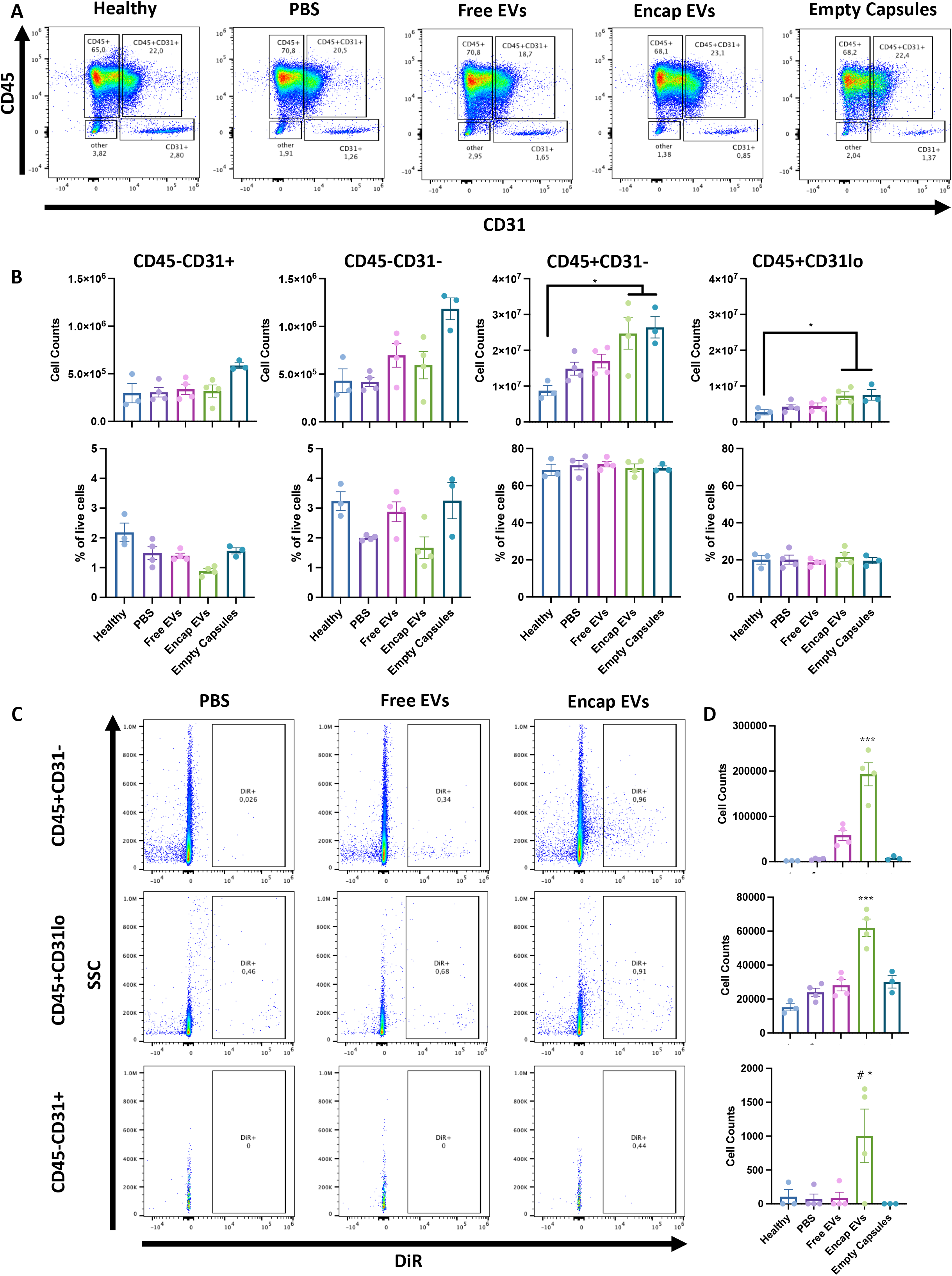
Local immune cell EV uptake increased by microencapsulation. Lungs were isolated, digested, and prepared for flow cytometry analysis 24h after DiR labelled EV administration. (A) Representative flow plots demonstrate consistent proportions of lung endothelial (CD45-CD31+), immune (CD45+CD31- or CD31lo), and other (CD45-CD31-) populations, which was quantified as cell counts, and proportions (B) demonstrating increased numbers of immune cells with the presence of microgels. EV uptake was assessed by evaluation of DiR labelling, representative flow plots are shown (C) and quantification (D) demonstrates significant EV uptake by CD45+CD31- and CD45+CD31lo cells when exposed to encapsulated EVs. Data represented as mean ± SEM, * p<0.05 PBS, Free EVs, and Empty Capsules compared to Encap. EVs, *** p<0.0005 all Tx compared to Encap EVs, # p = 0.07 healthy compared to Encap EVs.

## Discussion

Systemic delivery of free EVs is a promising treatment for a variety of lung diseases^33^, including PAH.^5^ However, after intravenous administration, EVs are rapidly cleared from the circulation, primarily by the liver and spleen,^15,17,34,35^ representing a significant challenge for therapies requiring targeted delivery of EVs to other tissues and organs, such as the lungs. Therefore strategies to improve organ specific EV retention and prolong the opportunity for EVs to interact with target host cells could greatly enhance efficacy of EV therapies. In this study, we demonstrated that encapsulation of MSC-EVs in nano-porous microgels significantly improved lung targeted EV retention and enhanced EV uptake by resident lung cells.

Microencapsulation has been previously employed for the delivery of therapeutic stem and progenitor cells.^22,23,27^ We sought to apply this technology for the targeted delivery of EVs to the lung. We demonstrated reduced uptake by macrophages and ECs of EVs from microgels *in vitro* over 24 hours suggesting a more sustained release system compared to free EVs which were readily taken up. This was also consistent with *in vivo* studies, which demonstrated excellent retention of microgels within the lungs up to a week following injection. In contrast, pharmacokinetic studies show that free EVs are rapidly cleared from the bloodstream within hours of administration^17^. Therefore, loading EVs into microgels represents a novel strategy to both enhance their delivery to the lung and increase their retention.

While previous studies have evaluated the overall biodistribution of systemically administered EVs, they have generally not examined which cells are responsible for EV uptake. To better understand the mechanisms by which EVs may exert their therapeutic actions, it is necessary to define the specific cell types with which they interact. In this study, we used fluorescently labelled EVs in combination with lung digestion and flow cytometry to gain insights into the cellular uptake of MSC-EVs within the lung. Loading EVs into microgels resulted in a marked increase in uptake by lung resident cells, primarily CD45+ immune cells. Uptake by lung resident immune cells could enhance the therapeutic efficacy of EVs in inflammatory lung diseases, such as acute respiratory distress syndrome. As well, in PAH it is hypothesized that MSC-derived EVs modulate the local inflammatory environment by interacting with immune cells, specifically macrophages^4^. While free EVs were readily taken up by macrophages *in vitro* (**Figure 1**), due to the short residence time *in vivo*, their ability to interact with lung host cells was limited resulting in minimal uptake by CD45+ immune cells after systemic delivery (**Figure 4**). In contrast, microencapsulation, which reduced *in vitro* uptake by MØ and ECs, greatly enhanced uptake *in vivo* by prolonging the retention of EV-loaded microgels in the lung and resulting in more sustained release kinetics after intravenous administration. However, whether the more efficient lung resident cell uptake of encapsulated EVs translates into greater therapeutic benefits remains to be seen.

The biomaterials used for microgel preparation will impact the tissue residence time and the local microenvironment. An agarose hydrogel formulation was developed for previous studies in our group designed to enhance cell persistence and engraftment following transplantation,^22^ (**Supplemental Figure 7**). This agarose-gelatin formulation was also effective in enhancing local retention of encapsulated EVs in the lung, yet there are opportunities to optimize the biomaterial formulation going forward. Immune cells were the primary cells involved in the uptake of EVs from microgels and even empty agarose-gelatin microgels promoted inflammation likely due to biomaterial-immune cell interactions inciting a foreign body response (FBR). The size and shape of implanted biomaterials has been shown to influence the FBR, although in this study no materials below 100μm were studied.^36^ Novel biomaterials have been developed to reduce the FBR, for example to reduce the inflammatory response to encapsulated pancreatic islets after transplantation.^37^ Biomaterials can also be modified to optimize the release kinetics of EVs. Recently, it has been reported that translocation of EVs through biomaterials and extracellular matrices is linked to the matrix mechanical properties and the EV deformability,^38^ and this represents an opportunity to modify the matrix properties to control the diffusivity of the loaded EVs. Alternatively, introduction of matrix metalloproteinase degradable sites through click-chemistries^39^ could further refine the release profile and determine the local retention time by enhancing degradation of the microgels.

This study demonstrates that EV-loaded microgels can be used to enhance EV delivery to the lungs compared to administration of free EVs. Furthermore, EV-loaded microgels act as a sustained release system to increase the time for interaction between host cells and EVs. Lastly, we have demonstrated that increased local retention of EVs results in significantly higher host cell uptake. All of this suggest EV-loaded microgels may offer significant benefits to enhancing delivery of EV therapies.

## Author Contributions

NDC has contributed to all aspects of this work including conception, performing the experiments, data analysis and preparation of this manuscript. KR and YD were involved with all animal studies. KR contributed technical knowledge and helped perform the biodistribution and *in vivo* digestion flow cytometry experiments. ABB and MG contributed expertise for the operation and maintenance of the microfluidic encapsulation device. DWC contributed to conception of the encapsulation platform, and in interpretation of results. DJS was involved in the conception and design of experiments, data analysis and interpreting results, and drafting this manuscript. DJS is accountable for all aspects of this work.

