## Supplemental Figures + Legends for "Targeting extracellular vesicle delivery to the lungs by microgel encapsulation"

**Supplemental Figure Legends**

**Supplemental Figure 1: Mesenchymal stromal cell derived extracellular vesicle characterization.** (A) Size distribution measured by nanoparticle tracking analysis demonstrating EVs with a median size less than 200nm. (B) Protein to particle ratio were consistent between cell sources tested. (C) MSC derived EVs expressed canonical EV markers CD63, CD9, and TSG101 with absence of Golgi membrane protein GM-130 by western blot.

**Supplemental Figure 2: Bone marrow derived macrophage characterization.** (A) Representative images showing morphology of untreated macrophages (MØ), IL4 stimulated M2-like, and LPS+IFNg stimulated M1-like macrophages. (B) Representative flow plots showing MØ were positive for immune markers CD45 and CD11b while negative for CD34.

**Supplemental Figure 3: Gating strategy employed for analysis of lung digestion samples.** In brief, debris and counting beads were excluded by FSC-A and SSC-A. Doublets were discriminated with SSC-A vs SSC-H. Dead cells were excluded. CD45 and CD31 were used to separate immune cells (CD45+) from endothelial cells (CD31+) from other (stromal, and epithelial CD45-CD31-). Finally, a DiR gate was applied on each subset to characterize the uptake of DiR labelled EVs. All gates were set using both FMOs and unstained controls.

**Supplemental Figure 4: Human umbilical vein endothelial cells (HUVEC) internalized EVs.** Representative confocal images of HUVEC exposed to pkh26 labelled EVs in vitro. Z-stack and layered image demonstrate presence of EVs within cytosolic space of the HUVEC indicating internalization.

**Supplemental Figure 5: Macrophage morphology was unchanged by EV exposure.** Representative images of all macrophage groups (MØ, IL4, and LPS+IFNg) with or without exposure to EVs in culture.

**Supplemental Figure 6: Free and encapsulated EV uptake in liver, spleen, kidney, and heart.** (A) Representative images of IVIS data showing DiR labelled EV distribution within animals treated with encapsulated and free EVs. PBS vehicle treated animals served as a control for background fluorescence intensity. (B) Quantification of fluorescence EV signal using average radiant efficiency demonstrating high uptake of free EVs in both liver and spleen. Minimal accumulation of encapsulated or free EVs was observed in the kidney or heart.

**Supplemental Figure 7: In vivo microgel degradation. (A) Schematic of study design.** (B) Representative images of empty microgels made of 2% agarose, and 1%agarose-1%gelatin by vortex emulsion, with size distribution (C) showing no difference in average size between these hydrogel formulations. (D) Empty microgel injection had no impact on right ventricular systolic pressures (RVSP) within these animals. (E) Representative H&E stained images of lung histology of lungs receiving empty microgels at 1-, 3-, and 7- days post injection. Red arrows indicate presence of microgels with evidence of surrounding immune infiltrate. (F) Quantification of capsules per lung section demonstrating enhanced clearance of 1%agarose-1%gelatin microgels.

**Supplemental Figure 8: DiR labelled EV uptake by CD45-CD31- cells. (**A) Representative flow plots showing minimal uptake of DiR+ EVs by CD45-CD31- cells, and (B) quantification showing no difference between any of the treatment groups.

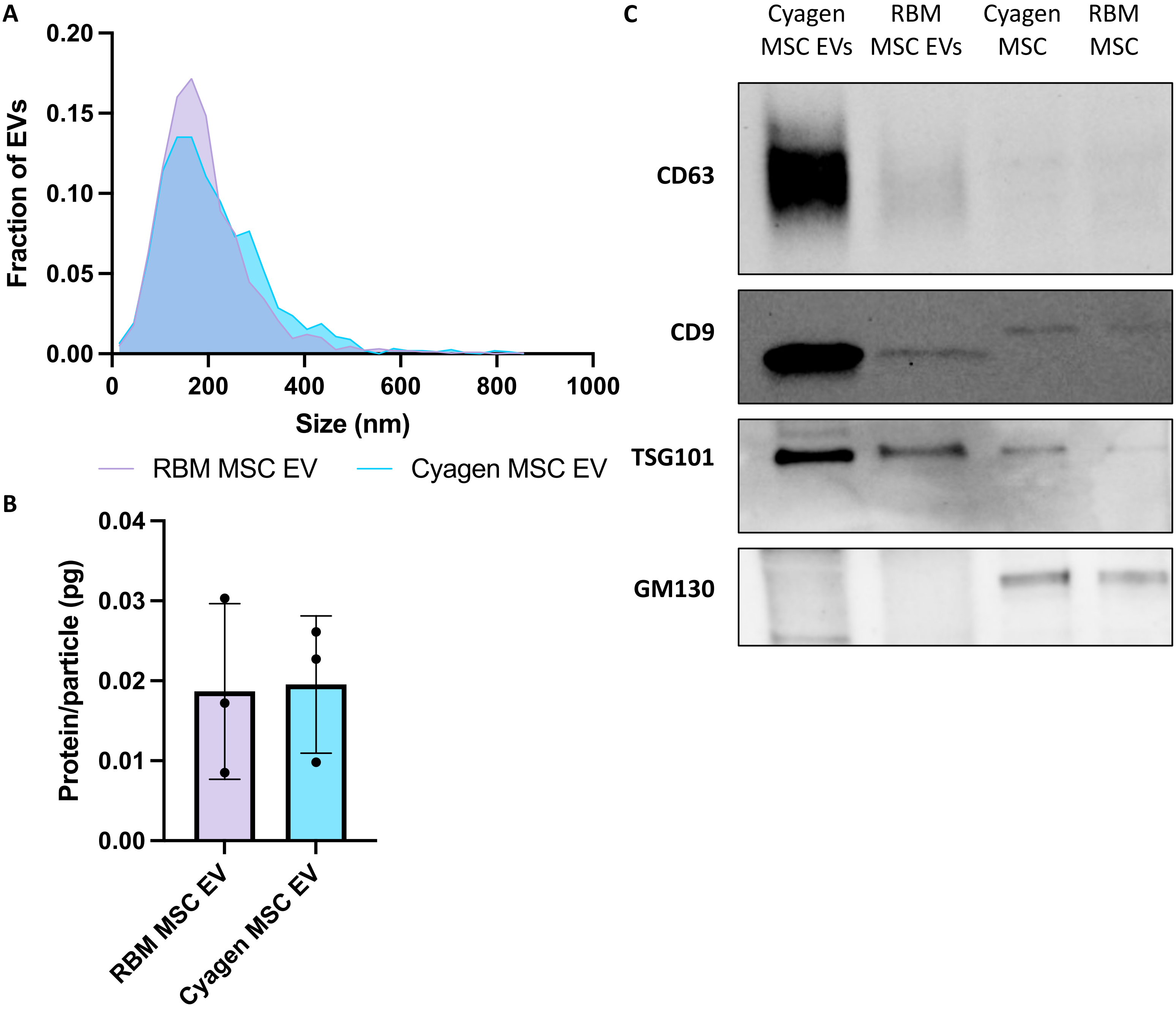

A
0.20
0.15
0.10
0.05
0.00
Fraction of EVs
0
200
400
600
800
1000
Size (nm)
RBM MSC EV
Cyagen MSC EV
B
0.04
0.03
0.02
0.01
0.00
Protein/particle (pg)
RBM MSC EV
Cyagen MSC EV
C
Cyagen MSC EVs
RBM MSC EVs
Cyagen MSC
RBM MSC
CD63
CD9
TSG101
GM130


Sup Fig 1

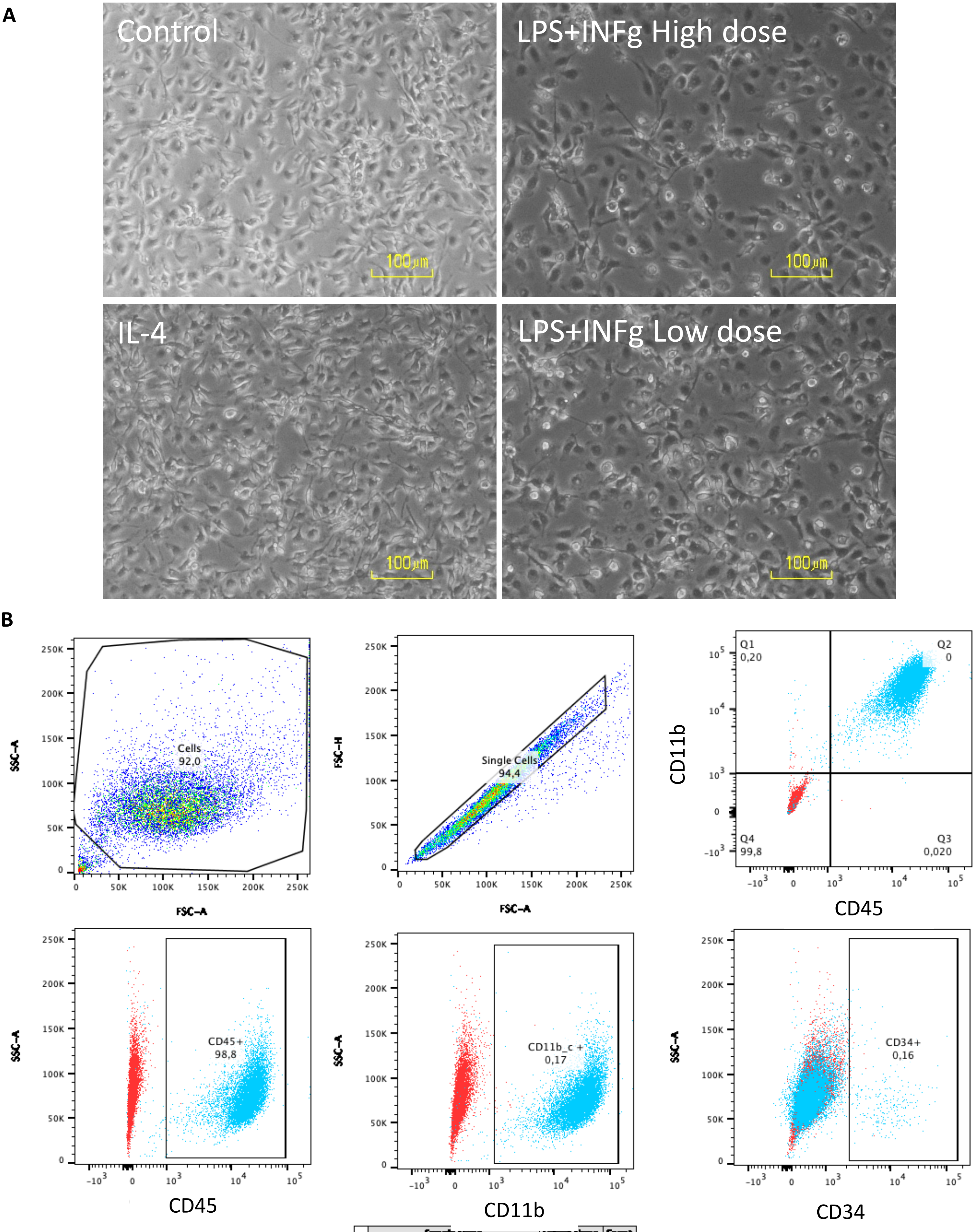
A
Control
LPS+INFg High dose
IL-4
LPS+INFg Low dose
100µm
B
Cells
92,0
SSC-A
FSC-A
Single Cells
94,4
FSC-H
FSC-A
Q1
0,20
Q2
0
Q4
99,8
Q3
0,020
CD11b
CD45
CD45+
98,8
SSC-A
CD45
CD11b_c +
0,17
SSC-A
CD11b
CD34+
0,16
SSC-A
CD34
Sample Name | Subset Name | Count
Specimen_001_Rat unstained_001.fcs | Single Cells | 9775
Specimen_001_Rat fully stained_002.fcs | Single Cells | 9503
Blue – Single Stain
Red – Unstained

Sup Fig 2

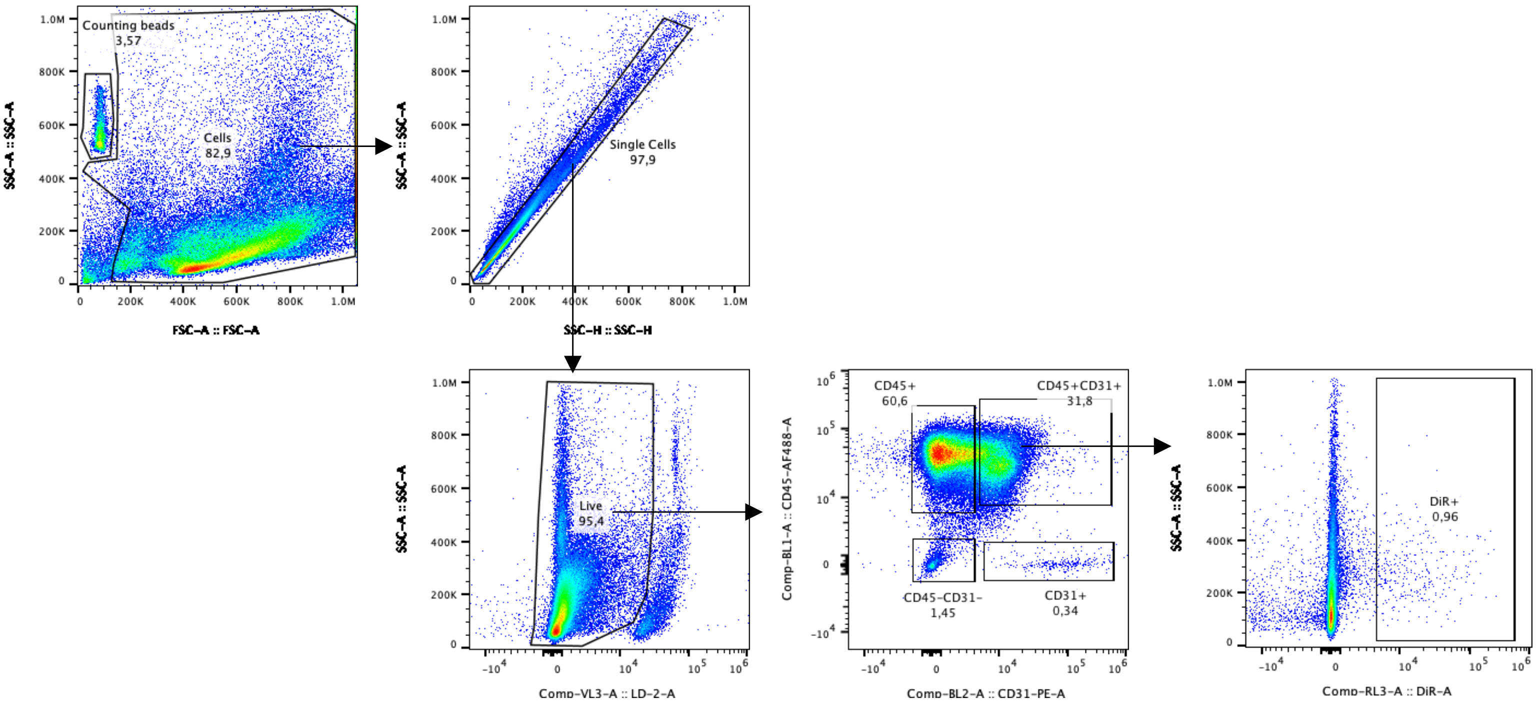
Counting beads
3,57
Cells
82,9
FSC-A :: FSC-A
SSC-A :: SSC-A
Single Cells
97,9
SSC-H :: SSC-H
Live
95,4
Comp-VL3-A :: LD-2-A
CD45+
60,6
CD45+CD31+
31,8
CD45-CD31-
1,45
CD31+
0,34
Comp-BL1-A :: CD45-AF488-A
Comp-BL2-A :: CD31-PE-A
DiR+
0,96
Comp-RL3-A :: DiR-A

Sup Fig 3

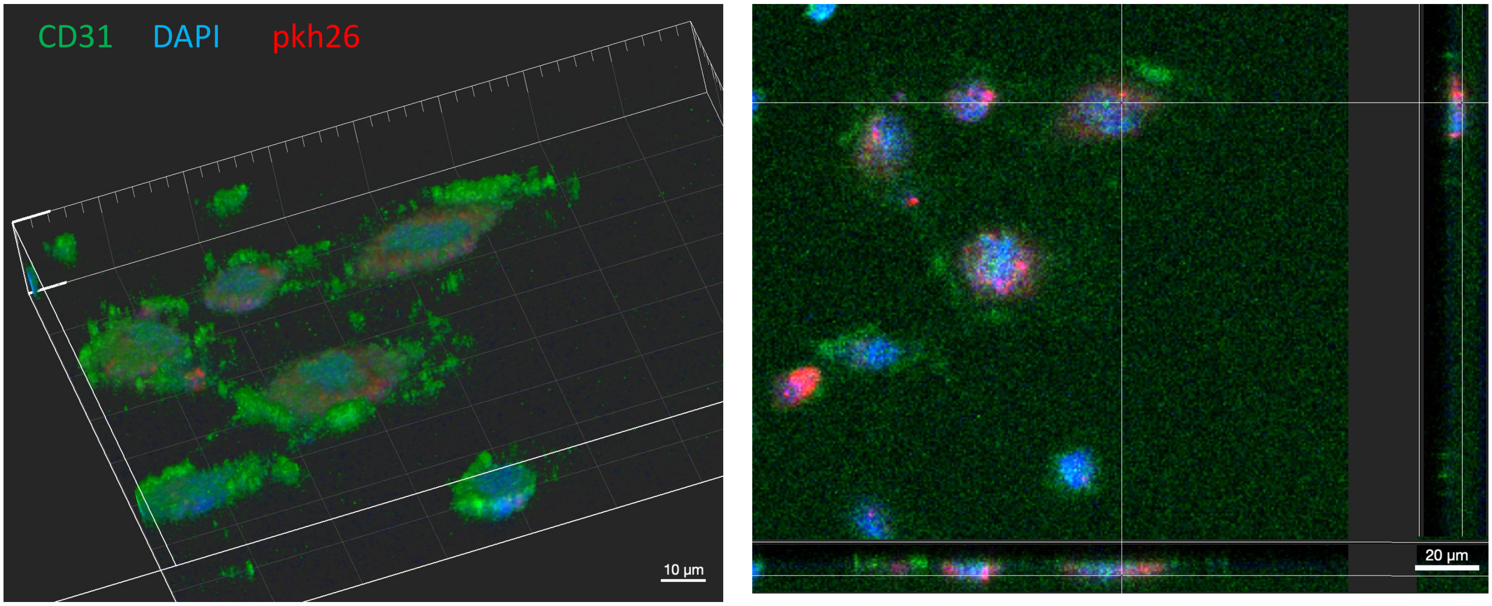


Sup Fig 4

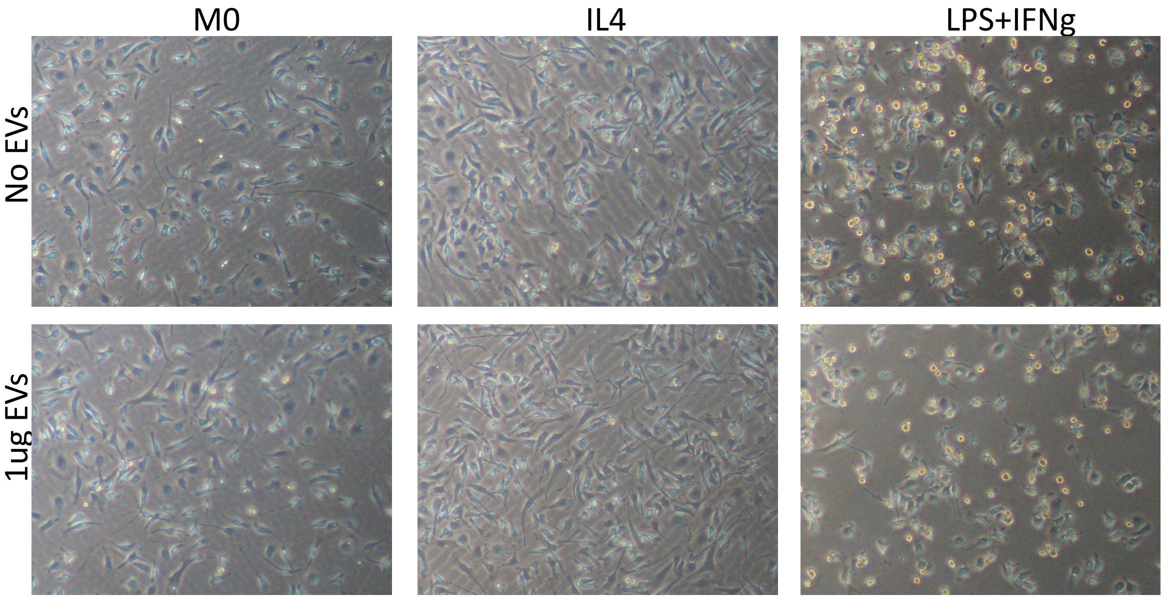


Sup Fig 5

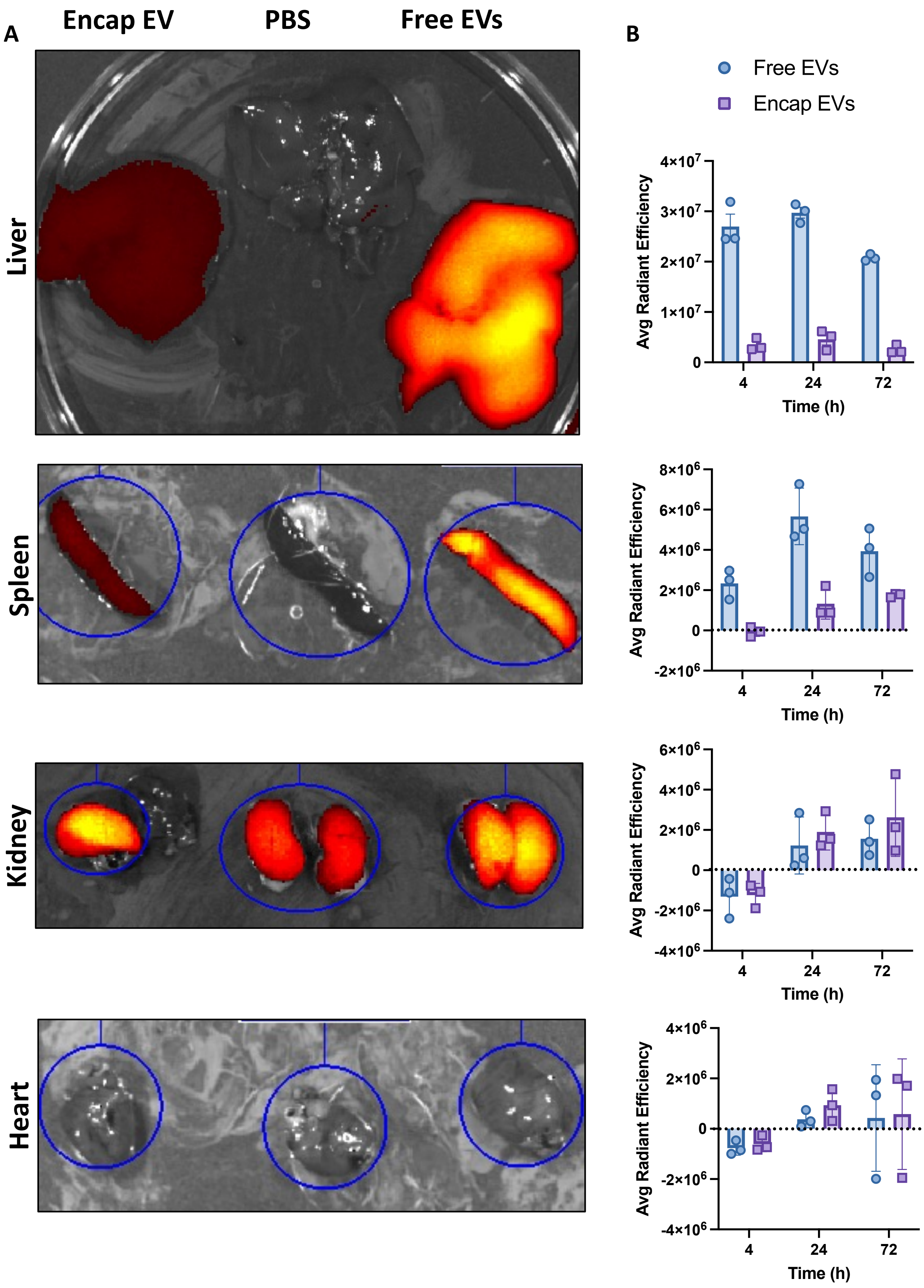
A
Encap EV
PBS
Free EVs
Liver
Spleen
Kidney
Heart
B
Free EVs
Encap EVs
Avg Radiant Efficiency
Time (h)
4
24
72
4×10^7
3×10^7
2×10^7
1×10^7
0
8×10^6
6×10^6
4×10^6
2×10^6
-2×10^6
-4×10^6

Sup Fig 6

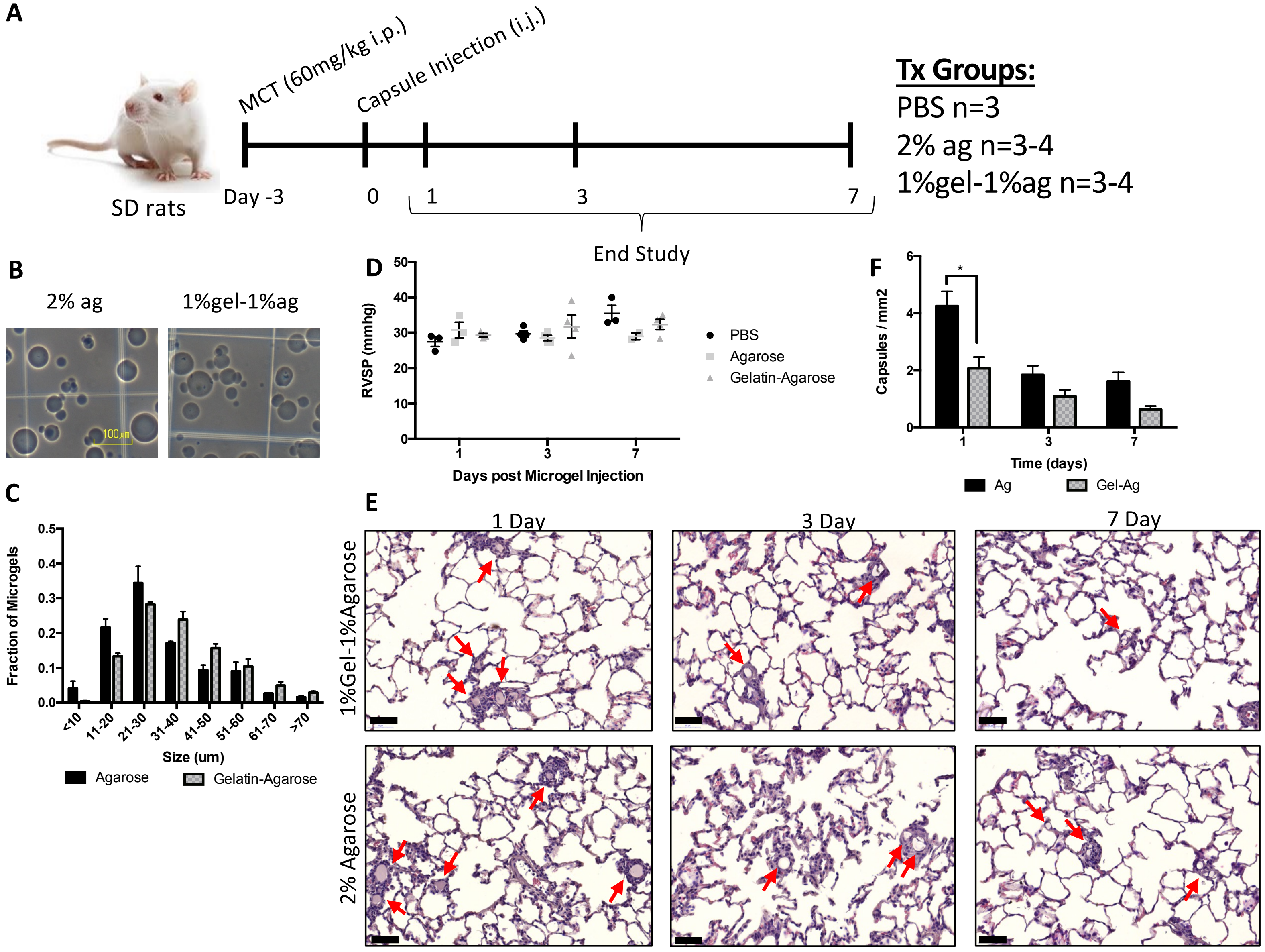
A
MCT (60mg/kg i.p.)
Capsule Injection (i.j.)
SD rats
Day -3
0
1
3
7
End Study
Tx Groups:
PBS n=3
2% ag n=3-4
1%gel-1%ag n=3-4
B
2% ag
1%gel-1%ag
100μm
C
Fraction of Microgels
Size (um)
Agarose
Gelatin-Agarose
D
RVSP (mmhg)
Days post Microgel Injection
PBS
Agarose
Gelatin-Agarose
F
Capsules / mm2
Time (days)
Ag
Gel-Ag
E
1 Day
3 Day
7 Day
1%Gel-1%Agarose
2% Agarose

Sup Fig 7

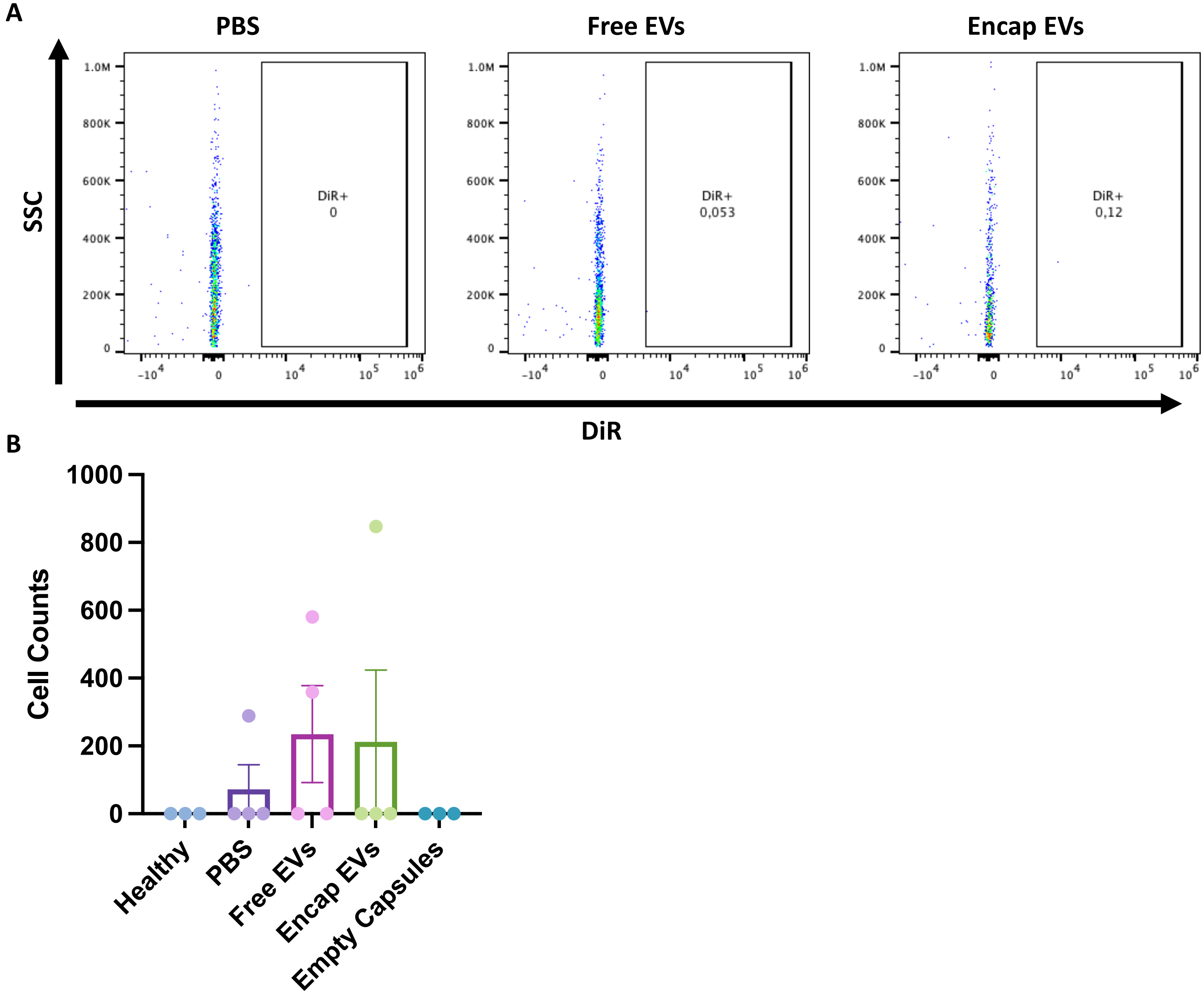

A
PBS
Free EVs
Encap EVs
SSC
DiR+
0
DiR+
0,053
DiR+
0,12
DiR
B
Cell Counts
Healthy
PBS
Free EVs
Encap EVs
Empty Capsules


Sup Fig 8
